## Supplemental Figures for "DNAJB1-PRKACA fusion protein-regulated LINC00473 promotes tumor growth and alters mitochondrial fitness in fibrolamellar carcinoma"

**A**

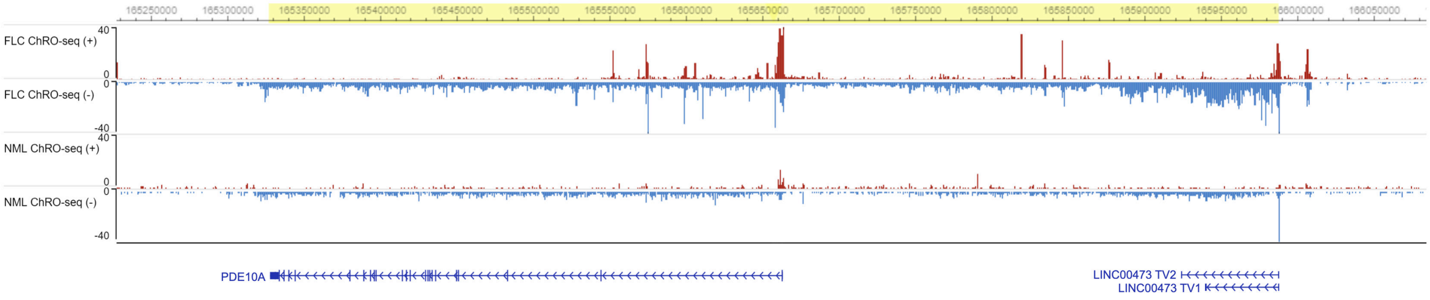


**B**

| Figure 1D | Figure 1E | Figure 1F |
|---|---|---|
| FCF06 | | |
| FCF09 | | |
| FCF26 | | |
| FCF27 | | |
| FCF34 | | |
| **FCF56** | **FCF56** | **FCF56** |
| **FCF83** | **FCF83** | **FCF83** |
| **FCF89** | **FCF84** | **FCF84** |
| **FCF90** | **FCF87** | **FCF87** |
| | **FCF89** | **FCF89** |
| | **FCF90** | **FCF90** |
| | **FCF82** | **FCF82** |
| | | FCF55 |
| | | FCF57 |
| | | FCF58 |
| | | FCF58L |
| | | FCF85 |

**C**

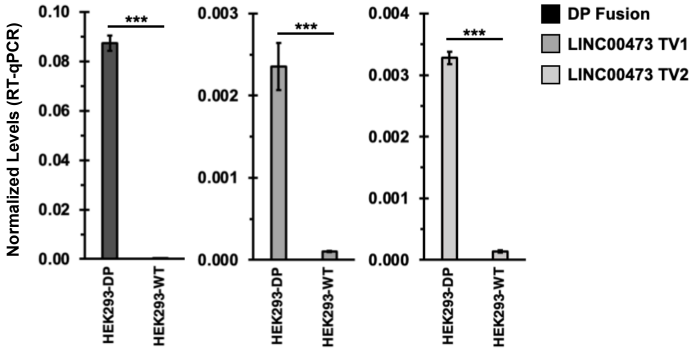


**Supplementary Figure 1. LINC00473 is a distinct transcription unit in FLC tumors.**

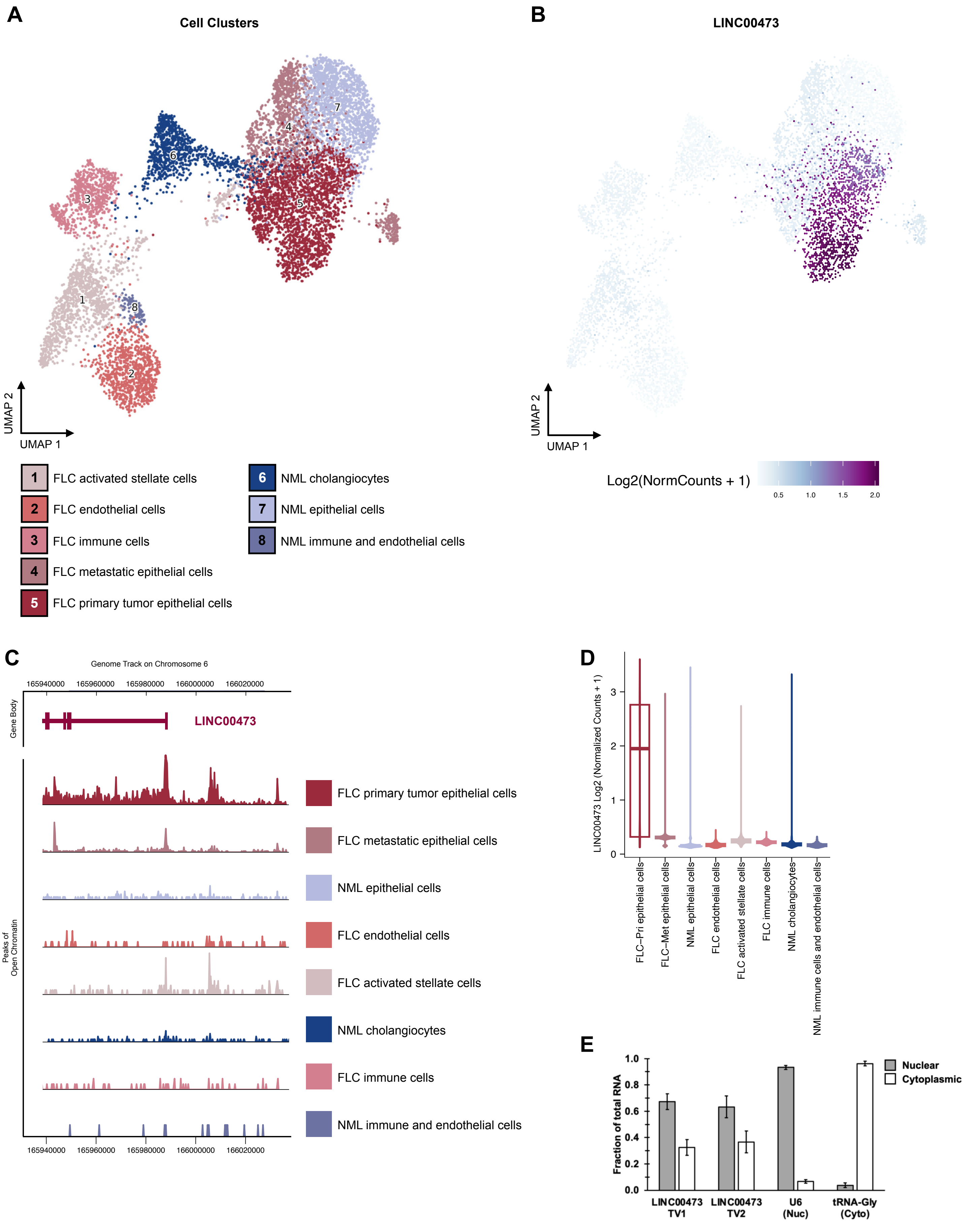


**Supplementary Figure 2. LINC00473 is enriched in FLC tumor epithelial cells.**

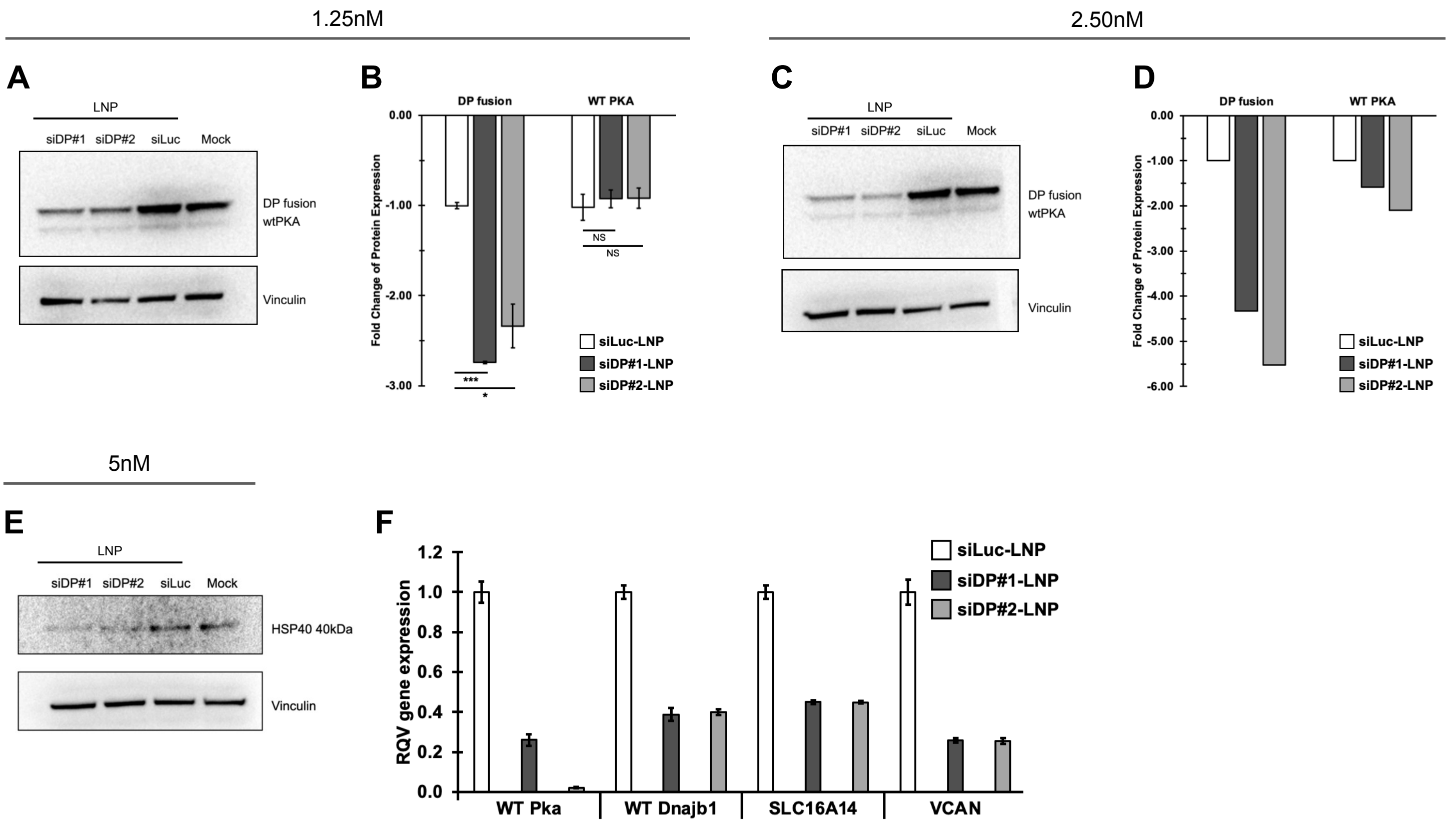


**Supplementary Figure 3. Silencing of the DP fusion.**

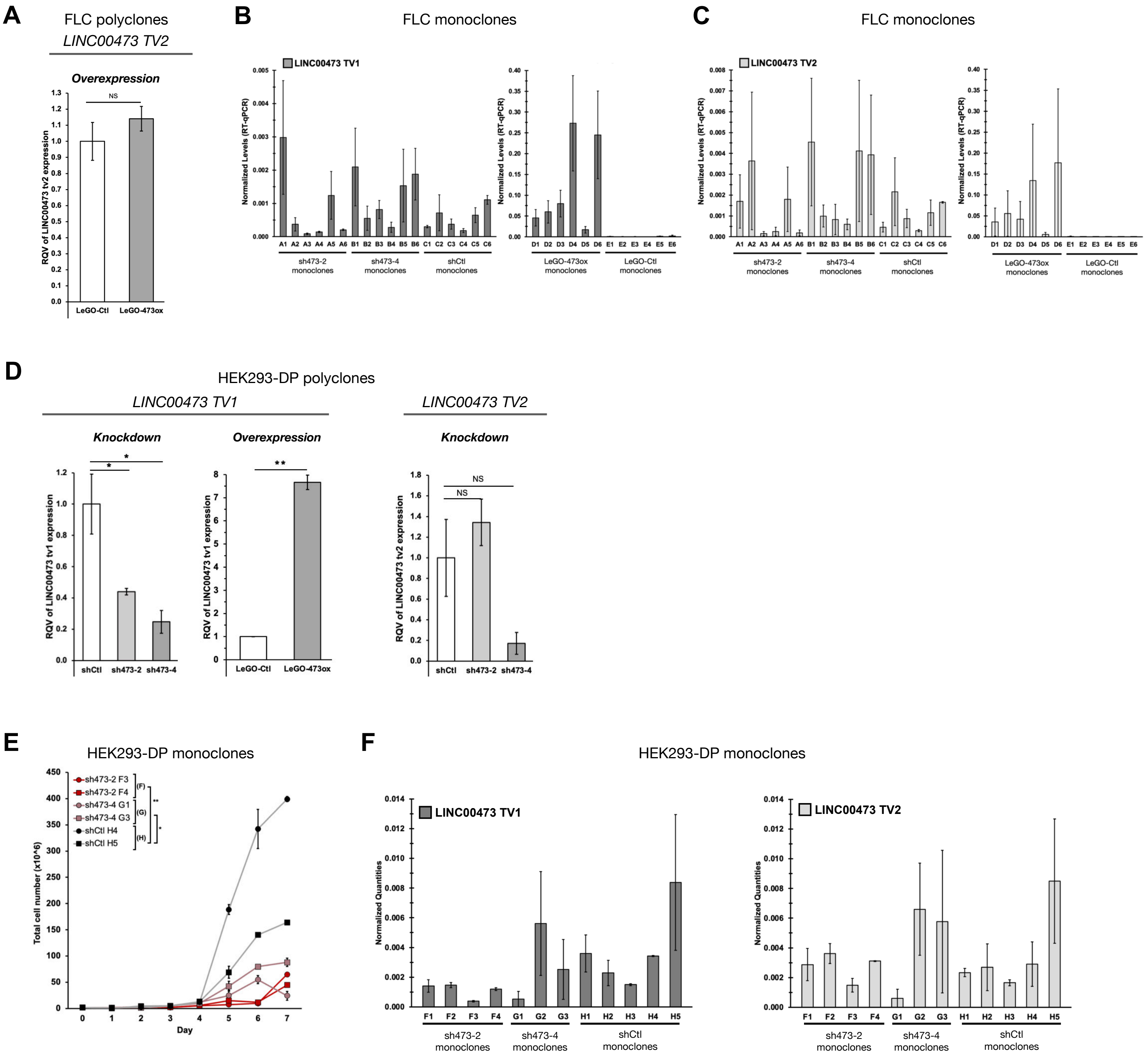


**Supplementary Figure 4. LINC00473 is efficiently downregulated or overexpressed in FLC and HEK293-DP cells.**

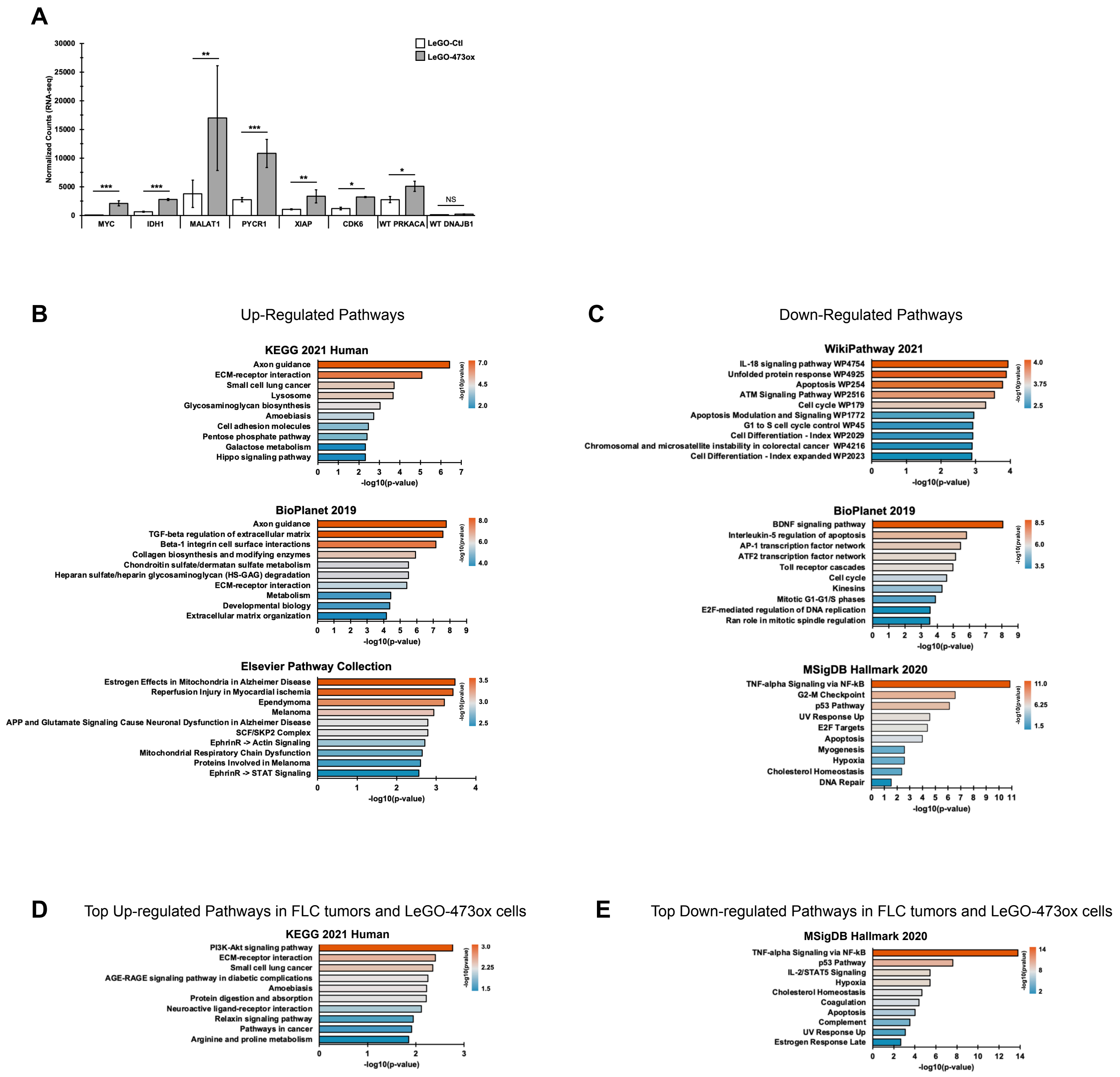


**Supplementary Figure 5. LINC00473 upregulates genes enriched in metabolism pathways and downregulates pathways related to apoptosis.**

**A**

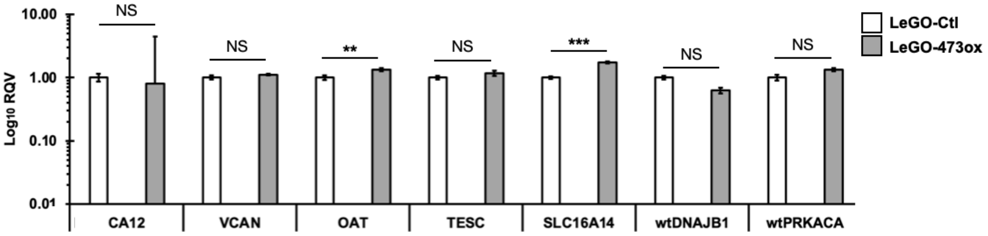


**Supplementary Figure 6. Gene expression in tumors generated from LINC00473 overexpressing cell implantation**.

**Supplementary Figure 1. LINC00473 is a distinct transcription unit in FLC tumors. (A)** Genome snapshot of the LINC00473 and *PDE10A* loci on chr6:165250000-166050000. Transcriptional signal on the plus and minus strand are shown in red and blue, respectively, in FLC tumors and non-malignant liver (NML) tissues. The RefSeq annotations for LINC00473 TV1 and TV2, and *PDE10A*, are shown on the bottom panel. Both genes are transcribed on the minus strand. LINC00473 has markedly higher levels of gene body transcription than the space in between LINC00473 and *PDE10A*. **(B)** Lists of primary patient tumors used in RNA-seq counts in a patient-matched subset from Cohort 1, gene expression via RT-qPCR in a patient-matched subset from Cohort 1, and correlation analyses on RNA expression levels of the DP fusion and LINC00473 isoforms TV1 and TV2 using patient tumors from Cohort 1, as described in Figures 1D, 1E, and 1F, respectively. Tumors indicated in red are common for all three analyses, and tumors highlighted in blue are shared for Figure 1E and 1F. **(C)** Normalized levels of DP fusion and LINC00473 isoforms TV1 and TV2 RNA in HEK293-DP cells relative to wild-type control (HEK293-Ctl). Normalized levels are 2^(-dCt) values using RPS9 for normalization and presented from 3 technical replicates. Data are represented as mean ± SD. P values are calculated by 2-tailed Student's t-test. $^*p < 0.05$, $^{**}p < 0.01$, $^{***}p < 0.001$.

**Supplementary Figure 2. LINC00473 is enriched in FLC tumor epithelial cells. (A)** snATAC-seq UMAP plot demonstrating eight cell clusters found in primary FLC, metastatic FLC, and NML tissue (~9500 nuclei). **(B)** Single-nucleus analysis of chromatin accessibility near the LINC00473 locus. Increasing signal is indicated by the color gradient (maximum signal is dark purple and minimal signal is light blue). **(C)** Top panel depicts the gene body of LINC00473, which is transcribed on the minus strand. Bottom panel demonstrates the quantification of snATAC-seq chromatin accessibility signal near the LINC00473 locus in different cell types. **(D)** Boxplot showing normalized counts of chromatin accessibility signal of LINC00473 in distinct cell types. **(E)** Subcellular fractionation followed by RT-qPCR in FLC cells using TaqMan primers designed to detect variants 1 (TV1) and 2 (TV2) of LINC00473, U6 (nuclear control) and tRNA-Gly (cytoplasmic control). Data represents n = 3 biological replicates ± SD. For A-D, data are represented from n = 1 FLC tumor, 1 Metastatic tumor and 1 NML.

**Supplementary Figure 3. Silencing of the DP fusion. (A, C)** Representative immunoblot of protein expression of DNAJB1-PRKACA (DP) fusion is detected with a protein kinase A catalytic α subunit (PKA) antibody. WT PKAc, DP fusion major, and DP fusion minor are identified. Lane 1, siDP#1-LNP; Lane 2, siDP#2-LNP; Lane 3, siLuciferase (siLuc-LNP) negative control; Lane 4, mock negative controls following 1.25nM treatment (A) or 2.50nM treatment (C) with siRNA-LNPs or mock condition overs 96 hours. Vinculin loading control is shown in the lower panel and run on the same blot**. (B, D)** Fold change of protein levels of the blot in panel A (B) and panel C (D), relative to siLuc negative control (n = 3). **(E)** Representative immunoblot of protein expression of WT DNAJB1. Lane 1, siDP#1-LNP; Lane 2, siDP#2-LNP; Lane 3, siLuciferase (siLuc-LNP) negative control; Lane 4, mock negative control. siRNA-LNP treatments at 5nM, or mock condition, over 96 hours. Vinculin loading control is shown in the lower panel and run on the same blot (n = 3)**. (F)** Gene expression from RT-qPCR following free uptake of siDP#1-LNP, siDP#2-LNP, and siLuc-LNP at 5nM treatment over 96 hours in FLC cells, as shown in Figure 3F (n = 3). Data are represented as mean ± SD. P values are calculated by 2-tailed Student's t-test. $^*p < 0.05$, $^{**}p < 0.01$, $^{***}p < 0.001$.

**Supplementary Figure 4. LINC00473 is efficiently downregulated or overexpressed in FLC and HEK293-DP cells. (A)** LINC00473 TV1 expression from RT-qPCR in FLC monoclonal cells following limiting dilutions of polyclonal cell lines to isolate single-clone cell colonies with stable gene knockdown using two independent shRNAs (sh473-2, sh473-4) or non-targeting shRNA control (shCtl) (n = 3), or overexpression using cDNA plasmid encoding LINC00473 TV1 (LeGO-473ox) or empty-vector control (LeGO-Ctl) (n = 3)**. (B)** Expression of LINC00473 TV2 was queried via RT-qPCR in FLC monoclones described in panel A (n = 3). **(C)** LINC00473 TV2 expression from RT-qPCR in FLC polyclones induced with LINC00473 TV1 overexpression (n = 3). **(D)** Expression of LINC00473 TV1 from RT-qPCR in HEK293-DP polyclonal cells following lentiviral transfection enabling stable gene knockdown using two independent shRNAs (sh473-2, sh473-4) or non-targeting shRNA control (shCtl) (n = 3) or overexpression using cDNA plasmid encoding LINC00473 TV1 (LeGO-473ox) or empty-vector control (LeGO-Ctl) (n = 3). The expression of second variant of LINC00473 (TV2) in HEK293-DP cells with stable sh473-2, sh473-4, or shCtl (n = 3)**. (E)** Cell growth curve of HEK293-DP monoclonal cells with stable LINC00473 knockdown (sh473-2: clones F3, F4; sh473-4: clones G1, G3) and non-targeting control (shCtl: clones H4, H5). Each monoclonal cell line was quantified 2 times across independent passages, and each point is the average cell count of 3 replicates. **(F)** Expression levels of both LINC00473 isoforms from RT-qPCR in HEK293-DP monoclones with stable gene knockdown using two independent shRNAs (sh473-2, sh473-4) or non-targeting shRNA control (shCtl). Data are represented as mean across 3 replicates ± SD**.** P values are calculated by 2-tailed Student's t-test. $^*p < 0.05$, $^{**}p < 0.01$, $^{***}p < 0.001$.

**Supplementary Figure 5. LINC00473 upregulates genes enriched in metabolism pathways and downregulates pathways related to apoptosis. (A)** Normalized counts of several genes of interest from RNA-seq of LeGO-473ox and LeGO-Ctl control FLC monoclonal cells. Bars are ± SD. The panel includes FLC-relevant transcriptional regulators (*MYC*), or genes associated with cellular metabolism (*IDH1*, *PYCR1*) and cell survival (*XIAP*, *CDK6*, *MALAT1*). **(B)** Pathway analyses using the 1403 upregulated genes in FLC monoclones with LINC00473 overexpression (LeGO-473ox) relative to empty vector control (LeGO-Ctl). Genes were filtered for expression with base mean > 100, log2FC > 1 and padj < 0.05 (DESeq). Pathways with adjusted p-value < 0.05 represented in figure. Color represents the -log10 adjusted p-value. **(C)** Pathway analyses using the 1374 downregulated genes in FLC monoclones with LeGO-473ox relative to LeGO-Ctl. Genes were filtered for expression with base mean > 100, log2FC < 1 and padj < 0.05 (DESeq). Color represents the -log10 adjusted p-value. **(D, E)** Gene list overlap analysis using significantly up- (D) or down-regulated (E) genes in FLC tumors relative to NML (n = 1497), and in LeGO-473ox cells relative to control (n = 1374). Pathways with adjusted p-value < 0.05 represented in figure. Color indicates the -log10 adjusted p-value. P values are calculated by 2-tailed Student's t-test. $^*p < 0.05$, $^{**}p < 0.01$, $^{***}p < 0.001$.

**Supplementary Figure 6. Gene expression in tumors generated from LINC00473 overexpressing cell implantation. (A)** Expression of FLC-related genes from RT-qPCR in LINC00473-overexpression (LeGO-473ox) and empty-vector control (LeGO-Ctl) subcutaneous FLC tumors. P values are calculated by 2-tailed Student's t-test. $*p < 0.05$, $**p < 0.01$, $***p < 0.001$.

| Type_Sample | Sample | Type | Seq source | Individual | Cancer Category | Sex | Age |
|---|---|---|---|---|---|---|---|
| FLC_FLC01_LNBS | FLC01_LNBS | FLC | UNC | FLC01 | Metastatic | F | 24 |
| FLC_FLC02_HMTK | FLC02_HMTK | FLC | UNC | FLC02 | Metastatic | M | 32 |
| FLC_FLC03_AAJH | FLC03_AAJH | FLC | UNC | FLC03 | Metastatic | M | 19 |
| FLC_FLC04_CDHM | FLC04_CDHM | FLC | UNC | FLC04 | Metastatic | M | 27 |
| NML_FLC06_SLVP | FLC06_SLVP | Normal | Cornell | FLC06 | Normal | F | 20 |
| FLC_FLC06_DZIS | FLC06_DZIS | FLC | Cornell | FLC06 | Metastatic | F | 20 |
| FLC_FLC07_LFOC | FLC07_LFOC | FLC | UNC | FLC07 | Primary | M | 16 |
| NML_FLC09_TZOG | FLC09_TZOG | Normal | Cornell | FLC09 | Normal | M | 28 |
| FLC_FLC09_MUOD | FLC09_MUOD | FLC | Cornell | FLC09 | Primary | M | 28 |
| FLC_FLC12_CTTF | FLC12_CTTF | FLC | Cornell | FLC12 | Recurrent | M | 27 |
| FLC_FLC13_QQEI | FLC13_QQEI | FLC | Cornell | FLC13 | Metastatic | F | 17 |
| FLC_FLC15_AWTJ | FLC15_AWTJ | FLC | Cornell | FLC15 | Metastatic | F | 48 |
| FLC_FLC17_WCKN | FLC17_WCKN | FLC | Cornell | FLC17 | Metastatic | F | 17 |
| FLC_FLC18_MKZC | FLC18_MKZC | FLC | Cornell | FLC18 | Primary | F | 18 |
| FLC_FLC20_ZDNV | FLC20_ZDNV | FLC | Cornell | FLC20 | Metastatic | M | 19 |
| FLC_FLC24_DJZW | FLC24_DJZW | FLC | Cornell | FLC24 | Primary | N/A | N/A |
| FLC_FLC25_UYHR | FLC25_UYHR | FLC | Cornell | FLC25 | Metastatic | F | 22 |
| NML_FLC26_ICBQ | FLC26_ICBQ | Normal | Cornell | FLC26 | Normal | N/A | N/A |
| FLC_FLC26_YJEE | FLC26_YJEE | FLC | Cornell | FLC26 | Primary | M | 18 |
| NML_FLC27_BWSX | FLC27_BWSX | Normal | Cornell | FLC27 | Normal | F | 18 |
| FLC_FLC27_BDCH | FLC27_BDCH | FLC | Cornell | FLC27 | Primary | F | 18 |
| FLC_FLC28_RKXK | FLC28_RKXK | FLC | Cornell | FLC28 | Metastatic | F | 30 |
| FLC_FLC29_QLXW | FLC29_QLXW | FLC | Cornell | FLC29 | Metastatic | F | 29 |
| FLC_FLC30_KPKS | FLC30_KPKS | FLC | Cornell | FLC30 | Metastatic | F | 31 |
| FLC_FLC31_OTOK | FLC31_OTOK | FLC | Cornell | FLC31 | Metastatic | F | 27 |
| FLC_FLC32_UOTX | FLC32_UOTX | FLC | Cornell | FLC32 | Metastatic | F | 16 |
| FLC_FLC33_NYTS | FLC33_NYTS | FLC | Cornell | FLC33 | Metastatic | M | 54 |
| NML_FLC34_PMVV | FLC34_PMVV | Normal | Cornell | FLC34 | Normal | M | 18 |
| FLC_FLC34_YROP | FLC34_YROP | FLC | Cornell | FLC34 | Primary | M | 18 |
| FLC_FLC55_T | FLC55_T | FLC | Cornell | FLC55 | Recurrent | F | 32 |
| NML_FLC56_N | FLC56_N | Normal | Cornell | FLC56 | Normal | F | 25 |
| FLC_FLC56_T | FLC56_T | FLC | Cornell | FLC56 | Primary | F | 25 |
| FLC_FLC57_T | FLC57_T | FLC | Cornell | FLC57 | Primary | F | 27 |
| FLC_FLC58L_T | FLC58L_T | FLC | Cornell | FLC58L | Primary | F | 27 |
| FLC_FLC82_T | FLC82_T | FLC | Cornell | FLC82 | Primary | M | 15 |
| NML_FLC83_N | FLC83_N | Normal | Cornell | FLC83 | Normal | F | 19 |
| FLC_FLC83_T | FLC83_T | FLC | Cornell | FLC83 | Primary | F | 19 |
| FLC_FLC84_T | FLC84_T | FLC | Cornell | FLC84 | Primary | M | 21 |
| FLC_FLC85_T | FLC85_T | FLC | Cornell | FLC85 | Metastatic | M | 12 |
| NML_FLC87_N | FLC87_N | Normal | Cornell | FLC87 | Normal | F | 31 |
| FLC_FLC88_T | FLC88_T | FLC | Cornell | FLC88 | Primary | F | 25 |
| NML_FLC89_N | FLC89_N | Normal | Cornell | FLC89 | Normal | M | 29 |
| FLC_FLC89_T | FLC89_T | FLC | Cornell | FLC89 | Primary | M | 29 |
| NML_FLC90_N | FLC90_N | Normal | Cornell | FLC90 | Normal | M | 36 |
| FLC_FLC90_T | FLC90_T | FLC | Cornell | FLC90 | Metastatic | M | 36 |
| FLC_FLC01_FHCC | FLC1 | FLC | FHCC | FLC01_FHCC | Primary | F | 14 |
| FLC_FLC02_FHCC | FLC2 | FLC | FHCC | FLC02_FHCC | Primary | F | 18 |
| FLC_FLC03_FHCC | FLC3 | FLC | FHCC | FLC03_FHCC | Primary | M | 14 |
| FLC_FLC04_FHCC | FLC4 | FLC | FHCC | FLC04_FHCC | Primary | F | 37 |
| FLC_FLC05_FHCC | FLC5 | FLC | FHCC | FLC05_FHCC | Primary | M | 23 |
| NML_FLC01_FHCC | NML1 | NML | FHCC | FLC01_FHCC | Normal | F | 14 |
| NML_FLC02_FHCC | NML2 | NML | FHCC | FLC02_FHCC | Normal | F | 18 |
| NML_FLC03_FHCC | NML3 | NML | FHCC | FLC03_FHCC | Normal | M | 14 |
| NML_FLC04_FHCC | NML4 | NML | FHCC | FLC04_FHCC | Normal | F | 37 |
| NML_FLC05_FHCC | NML5 | NML | FHCC | FLC05_FHCC | Normal | M | 23 |
